# An antibiotic-resistance conferring mutation in a neisserial porin: Structure, ion flux, and ampicillin binding

**DOI:** 10.1101/2020.11.06.369579

**Authors:** A. Bartsch, C.M. Ives, C. Kattner, F. Pein, M. Diehn, M. Tanabe, A. Munk, U. Zachariae, C. Steinem, S. Llabrés

## Abstract

Gram-negative bacteria cause the majority of highly drug-resistant bacterial infections. To cross the outer membrane of the complex Gram-negative cell envelope, antibiotics permeate through porins, trimeric channel proteins that enable the exchange of small polar molecules. Mutations in porins contribute to the development of drug-resistant phenotypes. In this work, we show that a single point mutation in the porin PorB from *Neisseria meningitidis*, the causative agent of bacterial meningitis, can strongly affect the binding and permeation of beta-lactam antibiotics. Using X-ray crystallography, high-resolution electrophysiology, atomistic biomolecular simulation, and liposome swelling experiments, we demonstrate differences in drug binding affinity, ion selectivity and drug permeability of PorB. Our work further reveals distinct interactions between the transversal electric field in the porin eyelet and the zwitterionic drugs, which manifest themselves under applied electric fields in electrophysiology and are altered by the mutation. These observations may apply more broadly to drug-porin interactions in other channels. Our results improve the molecular understanding of porin-based drug-resistance in Gram-negative bacteria.

## INTRODUCTION

As evidenced by the world-wide Covid19 pandemic, untreatable infectious diseases hugely impact on public health, the global economy and the lives of people across the world. The rise in previously-treatable infections becoming drug resistant, which is also currently experienced on a world-wide scale, poses a major problem for public health that urgently needs to be addressed [1]. Many drug-resistant infections are caused by Gram-negative bacteria, which exhibit a higher level of intrinsic resistance to antibiotics compared to Gram-positive bacteria owing to the complex architecture of their cell wall [2, 3]. The combination of insufficient inward drug permeation across the outer membrane - an additional lipid bilayer in the Gram-negative cell wall - and together with the activity of tri-partite efflux pumps spanning both the inner and outer membrane both contribute to poor drug susceptibility [4, 5].

Most antibiotic drugs permeate the outer membrane of Gram-negative bacteria through porins, beta-barrel proteins that form large water-filled channels in the membrane used by the organisms to import nutrients [6–8]. Down-regulation or mutations of porins, in conjunction with the upregulation of efflux pumps, has been reported to underpin drug-resistant phenotypes in Gram-negative bacteria [9, 10]. It is therefore important to understand the role of porin mutations in driving resistance to aid the design of improved antibiotic treatments.

The permeation of antibiotic molecules through porins is often studied by electrophysiology experiments. Voltage-induced ion flux across the channels can be partially or fully blocked upon drug binding or migration through the protein pore, revealing drug-channel interactions [11–13]. If binding events are of sufficient affinity, the structural details of such interactions can be determined by X-ray crystallography, while dynamic events such as transient association and drug migration are frequently studied in molecular detail using molecular dynamics simulations [14–16].

In this work, we used a combination of X-ray crystallography, experimental electrophysiology, and molecular dynamics simulations to study the role of mutations in PorB of *Neisseria meningitidis* – the causal agent of bacterial meningitis – on antibiotic binding and permeation. PorB is a trimeric anion-selective porin and the second most abundant protein in the outer membrane of *N. meningitidis*, possessing a binding site for antibiotics in its pore [13]. It plays an important role in the human immune response to neisserial infections and has also been linked to inducing apoptosis upon infection of host cells [17–19].

The only two pathogenic species of *Neisseriae* are *N. meningitidis* and *N. gonorrhoeae* [20]. Isolates of *Neisseriae* with intermediate resistance to antibiotics have been reported to exhibit single-point mutations in PorB [9, 21]. These mutations are located in the eyelet of the pore, the narrowest region within the channels, which contains a large number of charged amino acids acting as an electrostatic filter [22]. In *N. meningitidis* PorB, the equivalent variant to the known resistance mutation G120K from PorB from *N. gonorrhoeae*, where a glycine is replaced by a lysine on loop L3 (G103K), is position G103 [9, 21]. This residue is located within an extended loop structure between beta-strands 2 and 3 that folds back into the beta-barrel and shapes the eyelet region (Fig. 1).

**FIGURE 1.**
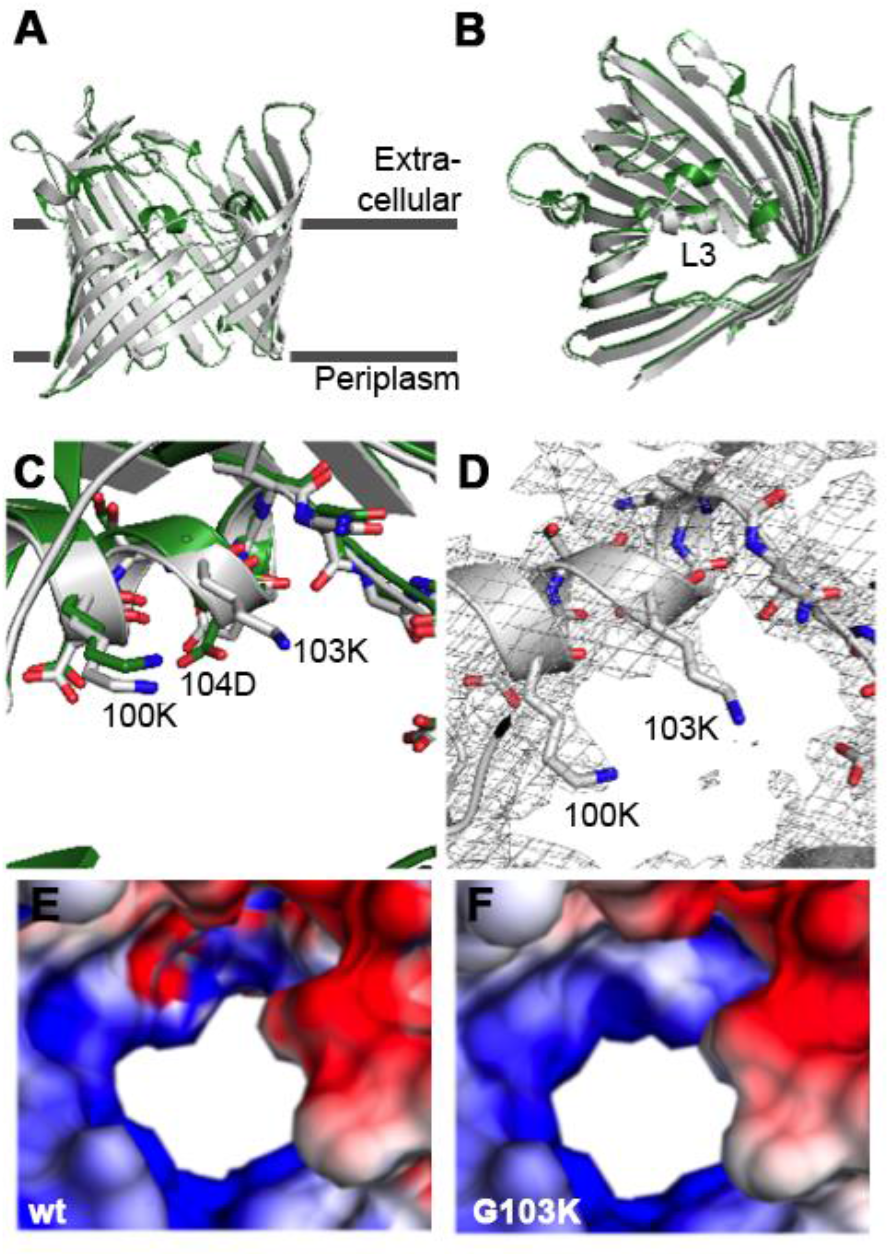
Overall structure of wt PorB from *N. meningitidis* (Nme) strain W135 (green) and PorB G103K (grey). (*A*) View from the membrane side; and (B) view from the extracellular side. (*C*) Representative electron density map for PorB G103K. The 2 *F*_o_-*F*_c_ density contoured at 1σ is shown. (*E, F*) Electrostatic surface potential of PorB wt (*E*) and one exemplar structure of PorB G103K (*F*) viewed from the extracellular side; surface potential calculated with APBS (positive potential in blue and negative potential in red, contoured from +8 *k*_B_*T*/*e* to −8 *k*_B_*T*/*e* at the solvent accessible surface area of PorB).

We recently characterised the binding interaction between wild-type PorB (wt PorB) and ampicillin through a combination of high-resolution electrophysiology recordings, molecular docking, biomolecular simulation and computational electrophysiology [13]. Here, we investigate the structural and functional effects of the resistance-related PorB mutation G103K and its interactions with ampicillin. Moreover, we observe distinct voltage-dependent effects on drug binding in the highly polarised eyelet of PorB, which are likely to apply more broadly to drug-porin interactions. The effect of the mutation on reducing ampicillin uptake is shown by liposome-swelling assays.

## RESULTS AND DISCUSSION

### Crystal structure and electrostatic surface potential of PorB

Crystals of G103K PorB from *N. meningitidis* W135 were grown under Jeffamine M-600 based conditions as previously reported [23, 24]. Since G103K PorB crystals were isomorphic to the previously determined crystals of wt PorB (PDB ID: 3VY8) [24], a simple refinement was performed to obtain initial model phases. As a result of manual model building and interactive refinement, G103K PorB has a final *R*_work_/*R*_free_ value of 21.3%/26.0% for data refined to 2.76 Å (Supporting Information).

The G103K single point mutation, located at the periplasmic side of loop L3, causes no significant changes in the secondary structure. The high degree of similarity between the G103K and wt PorB structures is reflected in a Cα rms deviation (RMSD) value of 0.278 Å (for 340 residues) (shown in Fig. 1A/B). Looking more closely at the mutation site, the loop L3 folds back from the extracellular side into the pore, forming an α-helix that constricts the pore to its narrowest point. The extracellular side of loop L3 slightly deforms the conformation of the α-helix, but this effect is also observed for wt PorB.

The electron density around lysine 103 was solved clearly from the α-to the δ-carbon atom, whereas the density of its positively charged ε-amino group could not be determined due to the potentially high mobility of the side chain (Fig. 1D). The bulkier side chain protrudes into the pore and decreases the open cross-section of the pore constriction, approximately by 3-5 Å as compared to wt, altering the PorB filter geometry.

For electrostatic analysis, first a model with 9 possible positions of the ε-amino group (out of 24) was constructed using the rotamer function in *Coot* [25], and the electrostatic surface potential was calculated for each structure using the Adaptive Poisson-Boltzmann Solver (APBS) method [26]. The remaining 15 positions of K103 Nε were excluded as potential conformations, because they were either located outside of the obtained electron density map, or due to steric hindrance with the side chains of N107, E128, K100 or the γ-carbon position.

Fig. 1 E/F shows the surface potential of wt PorB and one exemplar structure of the G103K mutant, viewed from the periplasmic and extracellular side. In wt PorB, opposite potentials are observed on each side of the eyelet. In contrast, the negative surface potential (red) completely disappears in G103K PorB, leaving a ring of positive potential around the eyelet. The positively charged constriction zone of PorB G103K is likely to influence the interaction of the porin with the β-lactam antibiotic ampicillin. We therefore conducted further investigations using electrophysiology and biomolecular simulations to characterise the interaction of ampicillin with the resistance-related G103K PorB mutant.

### Conductance of wild type and G103K PorB in planar lipid bilayer recordings

To determine the influence of the G103K point mutation on the conductance of the pore, we performed single channel measurements on wt and G103K PorB using solvent-free bilayers. Characteristic steps in the current signal were observed after addition of the protein (Fig. 2A/C), representing the opening and closing transitions (*gating*) of the channels.

**FIGURE 2.**
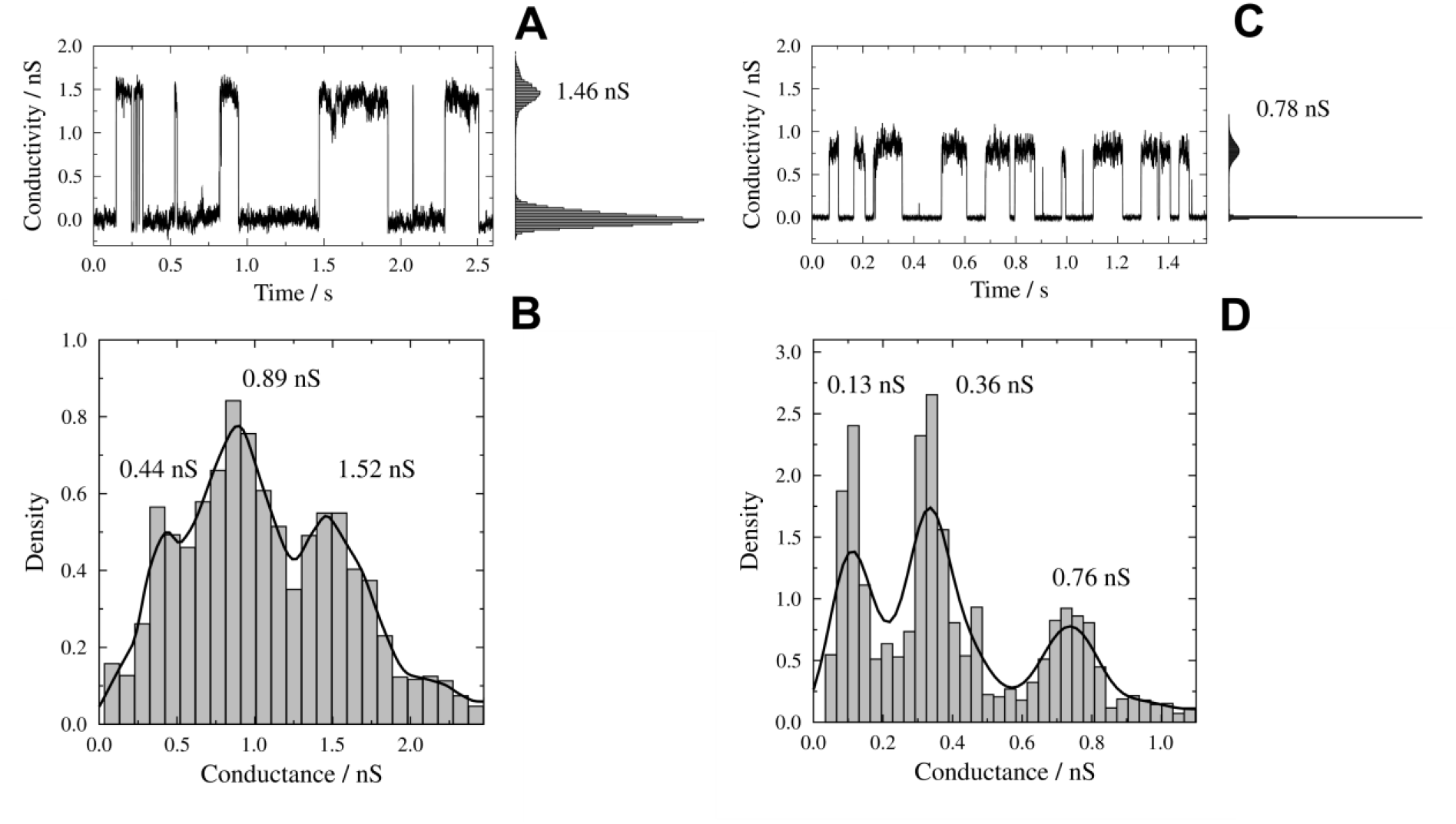
Comparison of the conductance states of wild type (wt) and G103K PorB. (*A*/*C*) Conductivity trace and corresponding point amplitude histogram of (*A*) PorB wt and (*C*) PorB G103K in a solvent-free membrane (DPhPC/Chol. 9:1) at *U* = +40 mV. (*B*/*D*) Conductivity event histogram including kernel density estimations using a Gaussian kernel (black solid lines) for (*B*) PorB wt (*G*_M,wt_ = (0.44 ± 0.18) nS, *G*_D,wt_ = (0.89 ± 0.26) nS and *G*_T,wt_ = (1.52 ± 0.46) nS) and (*D*) PorB G103K (*G*_M,G103K_ = (0.13 ± 0.05) nS, *G*_D,G103K_ = (0.36 ± 0.07) nS and *G*_T,G103K_ = (0.76 ± 0.12) nS). Buffer: 10 mM HEPES, 1 M KCl, pH 7.5. Bandwidths of 0.08 (*B*) and 0.05 (*D*) were chosen by visual inspection. Note that different bandwidths have been chosen to adjust for the different sizes of the conductance levels.

The conductance values of all gating events were included in a histogram and a kernel density was estimated using a Gaussian kernel (Fig. 2B/D). The evaluation resulted in three conductance states for both, wt and mutant G103K PorB, which can be assigned to the open state of a monomer (*G*_M_), dimer (*G*_D_) and trimer (*G*_T_) each. Compared to the conductance states of wt PorB, the conductance values for G103K PorB are significantly smaller with values of *G*_M,G103K_ = (0.13 ± 0.05) nS, *G*_D,G103K_ = (0.36 ± 0.07) nS and *G*_T,G103K_ = (0.76 ± 0.12) nS. The smallest conductance of 0.13 nS approaches a range where general current fluctuations of the membrane disturbed by the detergent are observed (Fig. S3).

### Pathways of ion transfer in PorB

We next conducted all-atom unbiased molecular dynamics simulations to investigate ion transfer pathways through the two studied channel variants. Both wt and G103K PorB trimers were embedded in 1-palmitoyl-2-oleoyl-*sn*-glycero-3-phosphocholine (POPC) model lipid bilayers surrounded by explicit water molecules. In accordance with previous computational studies [24, 27], wt PorB shows two distinct pathways for the transfer of anions and cations (Fig. 3A). In the G103K variant, the bulkier and positively charged lysine sidechain extends into the pore, thereby occupying the pathway for cations and leading to its disruption (Fig. 3B).

**FIGURE 3.**
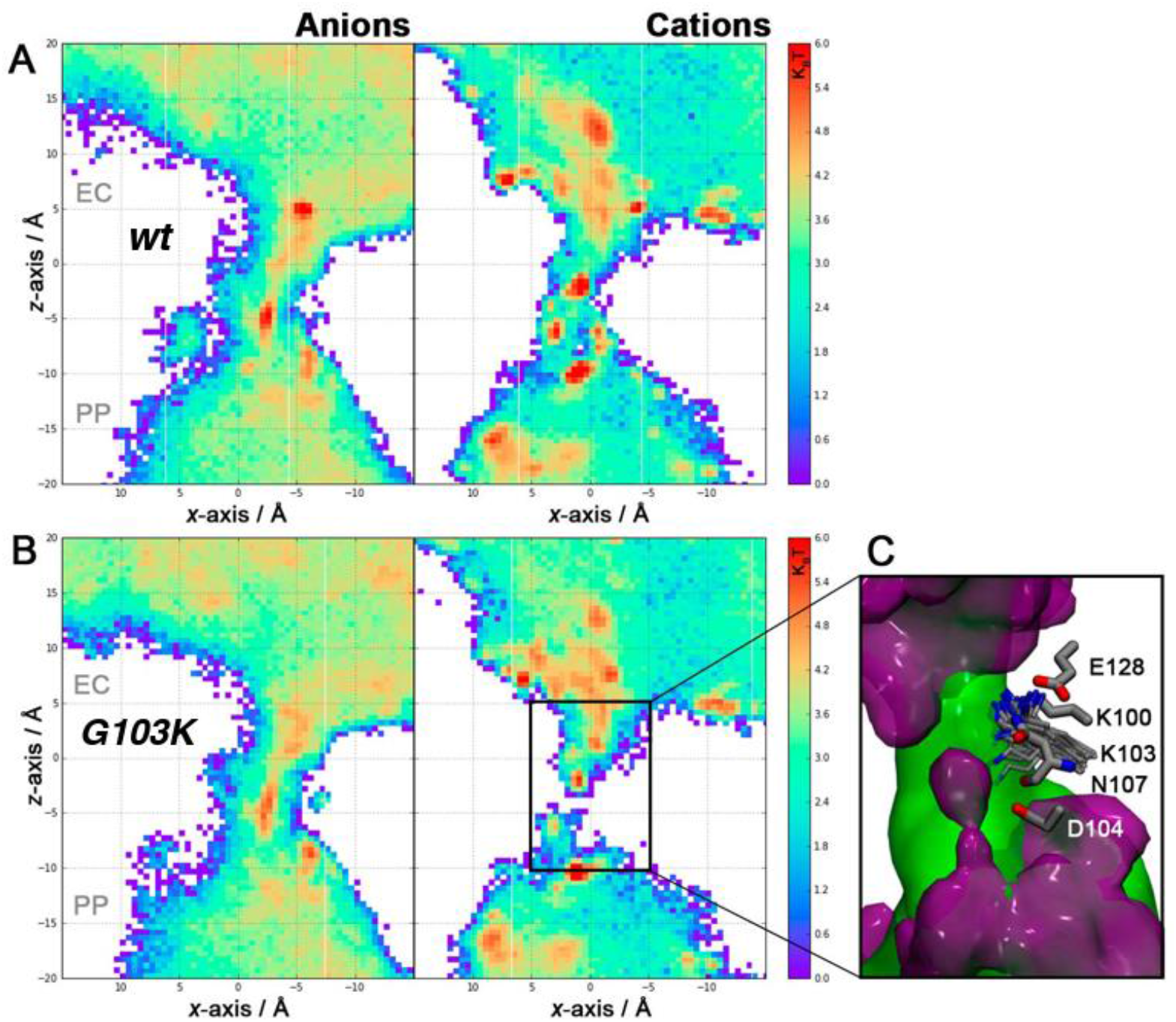
2D potential-of-mean-force free energy landscape (*k*_B_*T*) for K^+^ and Cl^-^ ions in the wt (*A*) and *(B)* G103K PorB channel. Both PMF shown are relative to the free energy minima. (*C*) Isodensity maps of cations (purple surface) and anions (green surface) in the interior of the G103K PorB channel. The isocontours correspond to an isodensity value of 0.015 particle Å^-3^.

The conformations of the K103 side chain sampled in our simulations are in good agreement with observations from the X-ray structure. Although the lysine residue can interact with the D104 side chain and the backbone of N108, it is mostly seen to either form a salt bridge with E128 or interact with passing chloride ions as shown in Fig. 3C. These interactions (Fig. S4) keep the lysine sidechain in conformations that protrude into the pore and therefore narrow the PorB channel.

Our experimental and computational electrophysiological data demonstrate that the conductance of G103K PorB is lower than that of the wt protein. These findings are in good agreement with results previously described for PorB from *Neisseria gonorrhoeae* (*Ngo*) [21]. Mutants of PorB (*Ngo*), including the mutant G120K, equivalent to the mutant G103K in PorB (*Nme*), exhibited smaller conductance states compared to the wild type of PorB (*Ngo*) [21].

### Ampicillin interaction with PorB

Binding of small molecules to the pore can result in short blockages of the open channel conductance if the ion flux is interrupted [11, 13, 28]. Traces of G103K PorB recorded after addition of ampicillin (Fig. 4A/B) show short interruptions of the open channel conductance which are not observed in its absence, indicative of the transient binding of the β-lactam antibiotic to the channel. The blockage events were shorter than 30 µs, which is below the time resolution given by the filter frequency of 5 kHz [13]. The filter frequency was chosen as the optimal compromise between noise reduction and signal smoothing, allowing us to evaluate residence times below 30 µs by using the analysis routine JULES together with a correction for missed events [29].

**FIGURE 4.**
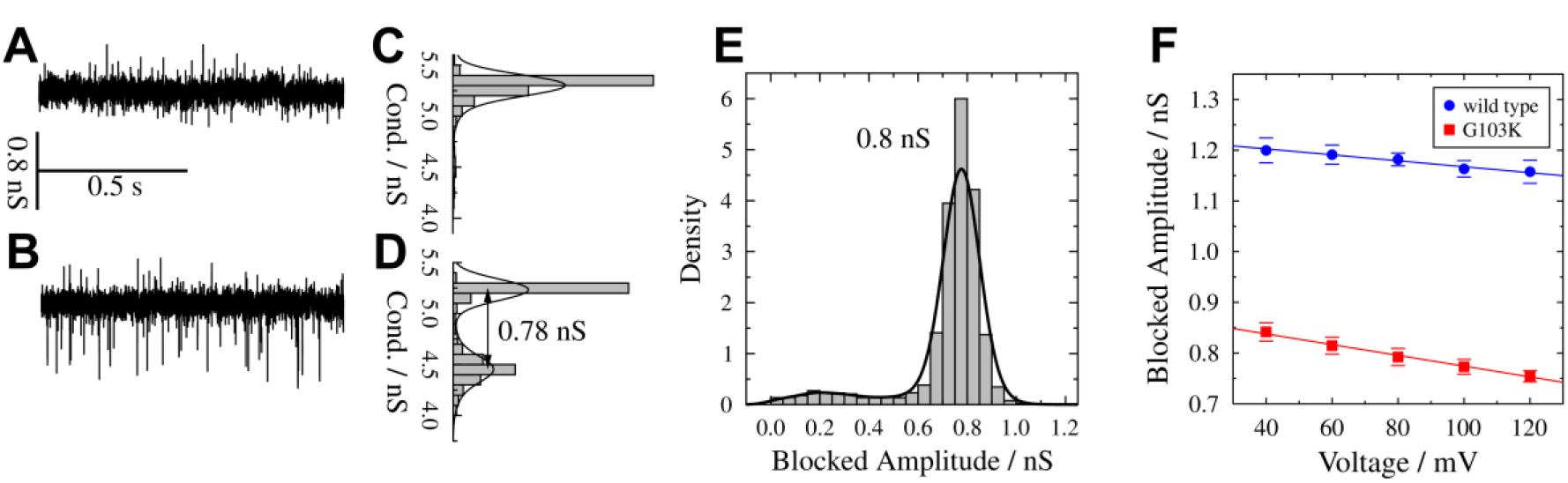
Electrophysiological recordings on G103K PorB in the presence of ampicillin. (*A-D*) Conductivity traces of G103K PorB at *U* = +80 mV in the absence *(A)*, and (*B*) in the presence of 1 mM ampicillin with the corresponding histograms of all conductance levels (*C,D*). The difference between the conductance levels after ampicillin addition is the *blocked amplitude G*_B_ plotted in (*E*) as an event histogram to determine the mean amplitude by Gaussian fits (black solid lines). (*F*) Blocked amplitudes for 1 mM ampicillin as a function of applied voltage for PorB wt (blue) and G103K (red). Four values were averaged; error bars show the standard deviation. For both proteins, a decrease of the amplitudes with increasing voltage is seen.

Histograms of all fitted conductance levels were generated (Fig. 4C/D), showing an additional conductance state in the presence of ampicillin (Fig. 4D). We assigned the new conductance state to the conductance of the blocked channel. The difference between the two states is a reduction in the open channel conductance due to ampicillin interaction, which we term the blocked amplitude *G*_B_. The corresponding event histogram is plotted in Fig. 4E with *G*_B,G103K_ = (0.8 ± 0.1) nS indicating that the entire trimer is blocked. The blocked amplitude is significantly lower for G103K PorB than for the wild type, in agreement with the larger conductance of the wild type trimer itself (Fig. 4F). However, in both cases a similar slight dependence on the applied voltage is observed.

We next determined the residence time and blocking frequency of ampicillin. The residence time is a characteristic feature of the interaction between an antibiotic and the pore. It reflects the time the drug molecule resides in the protein pore and blocks the flow of ions. If the rate constant of binding, *k*_on_, is unchanged, a longer residence time indicates a higher binding affinity. We analysed the residence time of wt and G103K PorB as a function of the ampicillin concentration in solution (Fig. 5A). As expected from other porin-antibiotics interaction studies [7, 30], the residence time is not influenced by the ampicillin concentration. A residence time of *τ*_R, wt_ = (35 ± 1) μs for wt PorB and a slightly larger one of *τ*_R, G103K_ = (44 ± 1) μs for G103K PorB was determined showing a stronger interaction of ampicillin with the mutant protein. Overall, the residence times are smaller than those found for other porins [7, 15, 31] indicating that the interaction is rather weak.

**FIGURE 5.**
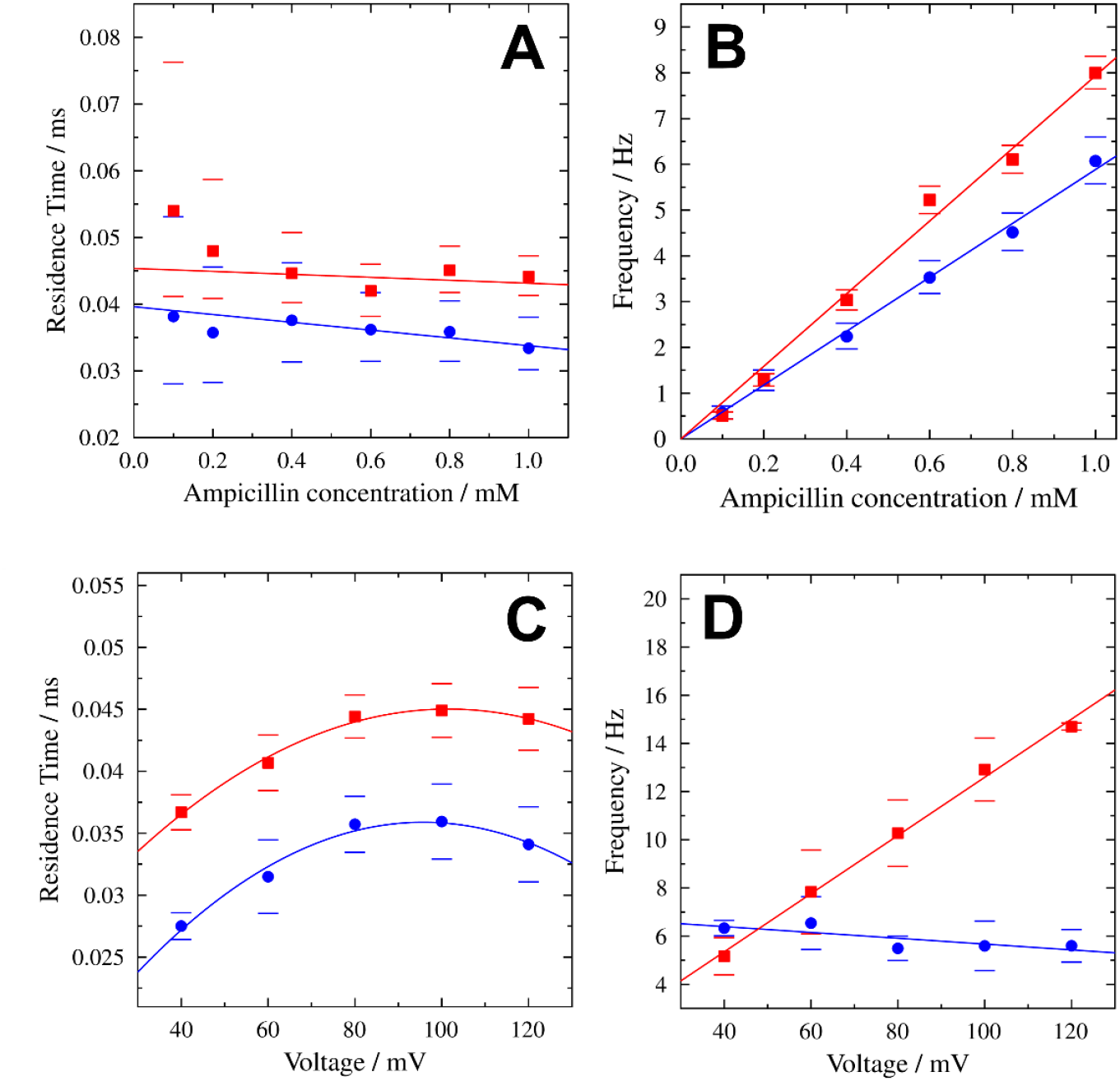
(*A*) Residence times and (*B*) blockage frequencies as a function of ampicillin concentrations for wt PorB (blue) and G103K PorB (red) including confidence intervals (bars) with 95% significance. The linear fit (weighted according to the confidence intervals) shows no significant dependence of the residence times on the ampicillin concentration. A residence time of *τ*_R, wt_ = (35 ± 1) μs for wt PorB and *τ*_R, G103K_ = (44 ± 1) μs for G103K PorB was determined using weighted means. For both proteins, the frequencies increase linearly with the ampicillin concentration. Recordings were performed at +80 mV. *(C)* Voltage-dependent residence times and (*D*) blockage frequencies of ampicillin for wt PorB (blue) and G103K PorB (red) in the presence of 1 mM ampicillin. The parabolic fits of the residence times serve as a guide-to-the-eye. Four measurements were averaged for each protein.

Both proteins show a bell-shaped dependence of the residence time on the applied voltage (Fig. 5B) with smaller residence times for the wild type protein in agreement with our previous findings (Fig. 5A) [13]. The voltage dependence of the residence time has been reported previously [7, 11, 15] and has been discussed in terms of an altered pore geometry induced by the applied voltage [32, 33].

A second characteristic of antibiotic binding is the frequency of blockage, i.e., how often the molecule interacts with the pore. The blockage frequency is expected to be linearly dependent on the concentration of the antibiotic as a result of the increasing number of available molecules [7, 11], which is confirmed by our measurements (Fig. 5B). The voltage dependence of the blocking frequency differs significantly between the two protein variants, increasing linearly with applied voltage for G103K PorB, whereas it decreases for wt PorB (Fig. 5D). This finding might be rationalized by the electric field within the pore, which is altered by the mutation at position 103. This can influence the orientation of the antibiotic as has been shown for mutated OmpC proteins [34].

### Ampicillin binding in wt and mutant PorB characterised by Computational Electrophysiology

To further understand the interaction between ampicillin and the mutated porin, we performed molecular docking calculations and all-atom unbiased molecular dynamics simulations of the PorB-ampicillin complex. Leveraging our recent work on ampicillin binding to the wt PorB channel, we used a similar molecular docking protocol employing two different docking software packages (GOLD[35] and rDOCK[36–38]) to find putative binding modes of ampicillin in the G103K pore. We identified a putative binding mode for ampicillin on the mutated PorB pore, which resembles the binding mode identified in the wt PorB channel [13] and other porins [14]. This binding mode was stable during 200 ns-long simulations of the complex for both wt and mutant PorB (Fig. S4 and Fig. S6).

Ampicillin is located at the extracellular vestibule of the PorB channel and largely obstructs the pore in both wt and G103K PorB channels. Ampicillin adopts a zwitterionic form at the studied pH (pH=6) and is therefore able to establish ionic interactions with both sides of the eyelet. The charged amino group of the antibiotic interacts with the E116 sidechain and the backbone oxygen atom of E110, while the carboxylic acid moiety binds to the sidechains of residues R130 and K77. In the G103K channel, the charged amino group of mutated residue K103 establishes an additional H-bond with the oxygen atom of the beta-lactam moiety.

In order to replicate the conditions of the electrophysiological experiments, we performed altogether eight 200 ns-long molecular dynamics simulations under voltage of both wt and G103K PorB channels using the computational electrophysiology (CompEL) method [27, 39]. The systems were duplicated along the *z*-axis and several ion imbalances were used to simulate positive and negative voltage ranges between ∼±120 – ∼±600 mV. Due to the double bilayer set-up of the system, this means that 16 PorB trimers were simulated in total. Conductance values of both PorB channels were calculated in the presence and absence of ampicillin, and we also tested its dependence on the direction and the strength of the applied electric field.

In the apo wt PorB channels, we observed a PorB trimer conductance of *G*_T,wt_comp_ = 1.54 ± 0.10 nS at positive voltages and *G*_T,wt_comp_ = 1.78 ± 0.14 nS at negative voltages. As reported previously, the values for the wt PorB channel are equal within their error margins and in agreement with the experimental values reported here and in previous computational studies. The flux of ions and water molecules is nearly symmetric at both negative and positive voltages as shown in other porins.

Upon ampicillin binding we observe a drastic reduction of the conductance of both porin variants as expected due to the partial blockage of the pore. The average wt PorB trimer conductance is *G*_T,wt,B,comp_ = 0.24 ± 0.04 nS at positive voltages and *G*_T,wt,B,comp_ = 0.69 ± 0.13 nS at negative voltages. Interestingly, the conductance is partially recovered at negative voltages as a consequence of the labile binding of the antibiotic, which is moved out of its binding site by pressure exerted by the force of the electroosmotic flow (EOF) arising due to the flux of hydrated ions under voltage. The EOF partially detaches the acidic side of the ampicillin in wt PorB at negative voltages as previously shown in Bartsch et al. [13]. This result is in good agreement with the experimental electrophysiology recordings, which show that ampicillin only blocks PorB at positive voltages.

We observe similar trends for the calculated conductance values in the G103K PorB variants. The G103K PorB trimer conductance is reduced from *G*_T,G103K,comp_ = 1.29 ± 0.07 nS at positive voltages and *G*_T,G103K,comp_ = 1.53 ± 0.09 nS at negative voltages in the apo form to *G*_T,G103K,B,comp_ = 0.09 ± 0.10 nS at positive voltages and *G*_T,G103K,B,comp_ = 0.30 ± 0.13 nS at negative voltages upon drug binding. The drop in conductance is asymmetric with respect to different polarisations of the membrane, which is consistent with a voltage-induced effect on ligand binding (see below). We also observe the partial unbinding of the beta-lactam antibiotic from the basic side of the eyelet. In mutant PorB, the flux of ions and water molecules is slightly reduced compared to the ampicillin bound wt PorB trimer, most likely related to the reduced size and altered electrostatic features of the pore.

### Ampicillin binding mode is field dependent

In our previous paper [13], we explained asymmetries seen in the ampicillin blockade of PorB current by the electro-osmotic effect (EOF). The EOF describes the stabilisation or destabilisation of ampicillin binding, depending on the polarisation of the membrane, due to water flow associated with the voltage-induced flux of hydrated ions across the porin eyelet [40, 41]. Stabilisation of ampicillin binding would lead to a longer and more frequent blockade, whereas destabilisation of binding would give rise to a partial recovery of the current. We also speculated that at higher voltages, the stronger electric field in the eyelet might result in a rotation of the zwitterionic ligand bound perpendicular to the field, yielding a further destabilisation of binding with increasing voltages. Experimentally, it can be shown that the frequency of ampicillin blockade decreases with increasing positive voltage in wt PorB, as opposed to the enhanced binding of neutral ligands under the influence of voltage-dependent EOF in alpha-hemolysin, while the frequency in G103K PorB follows the trend observed in alpha-hemolysin [40]. We therefore investigated the voltage dependence of ligand binding in both wt and G103K PorB.

In our present high voltage simulations (CompEL ion imbalance Δq = 18, Vm ≈ ±600 mV), the drug shows a rotation relative to the z-axis that depends on the voltage across the membrane (Fig. 6A/B). The acidic moiety moves in the direction of the positively polarised face of the membrane, while the basic moiety rotates towards the negative face. This leads to opposing rotations of the bound drug in its binding site under negative and positive voltages. In simulations of wt PorB without voltage (Δq = 0), no translation or rotation of ampicillin is observed (Fig. S6). Interestingly, the rotation of the drug is more pronounced in wt PorB than in the G103K variant (Fig 6B). In the mutant, voltage-dependent translation of the molecule slightly dominates over field-induced rotation (Fig. S8).

**FIGURE 6.**
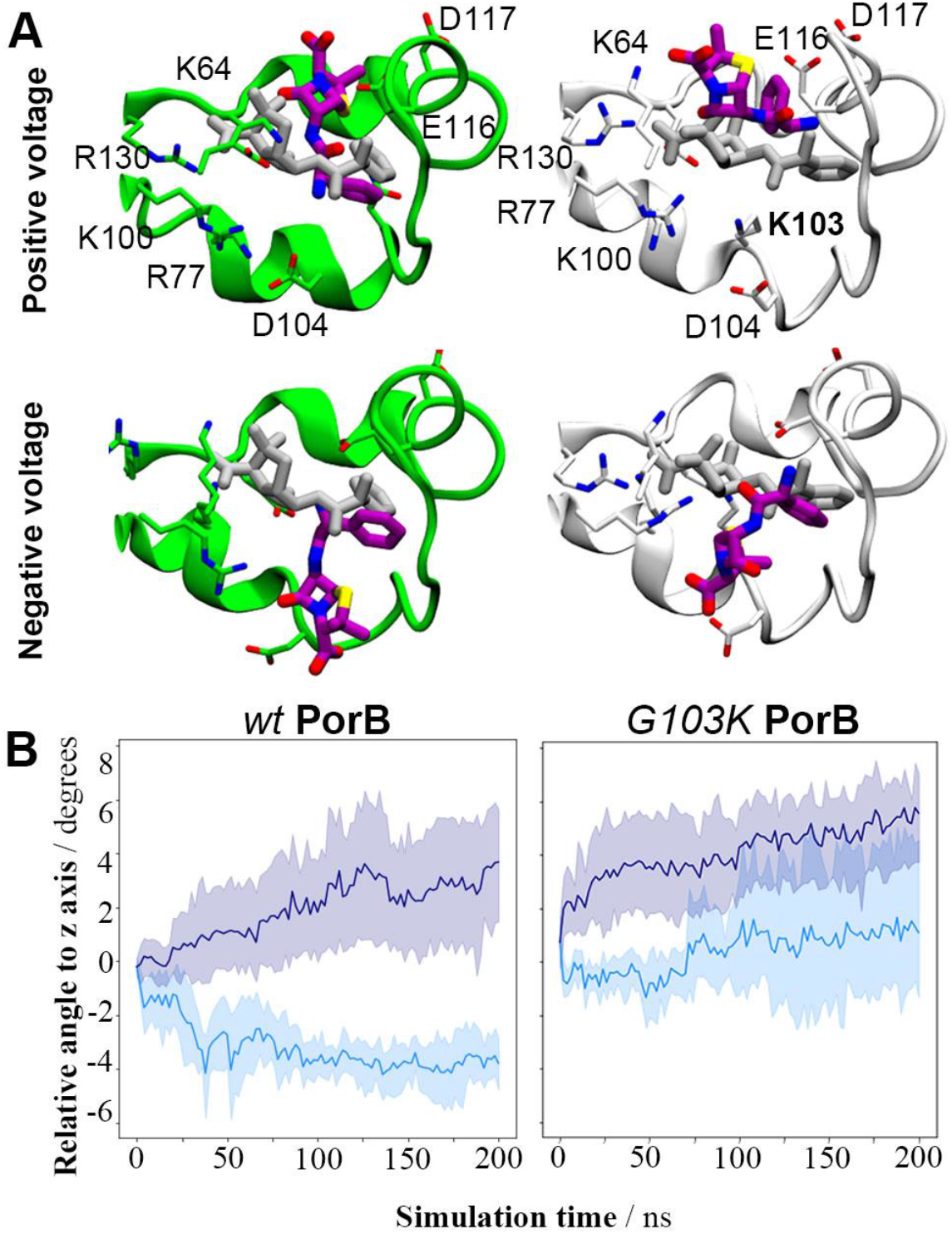
Comparison of field-dependent behaviour of ampicillin in wt and G103K PorB systems. *(A)* Final configurations of the ampicillin bound to both PorB proteins. Green and white cartoons represent the L3 loop of wt (PDB ID: 3VY8) and G103K PorB (PDB ID: 7DE8) proteins respectively. Initial and final orientation of ampicillin molecules are represented as grey and purple sticks, respectively. *(B)* Field-dependent rotation of ampicillin relative to the *z*-axis in wt (left) and G103K PorB (right). A greater difference between angles under positive and negative voltages emerges in wt PorB. The membranes under positive and negative voltages are depicted in navy and cyan, respectively, for a CompEL ion imbalance Dq of 18e (V_m_ ≈ ±600 mV). Each channel is treated individually with the mean value and a confidence interval of 95% plotted every 2 ns.).

The EOF exerts pressure on a molecule bound to a pore along the direction of ion flux, i.e. usually the channel axis. This leads to the expectation that voltage-dependent translation out of the binding site will be observed if the EOF predominates, while rotation of a zwitterion bound perpendicular to the membrane electric field is likely to be a direct result of the applied voltage. It appears that the mutation leads to a switch from a mainly direct effect caused by the field at high voltages, rotating ampicillin in its binding site in wt PorB, to increasing EOF-induced translation in G103K PorB. We suggest that the difference seen experimentally for the voltage-dependent frequency of pore blockade between the wt and the mutant (Fig. 5D) can be explained by this switchover of predominant effects. The mutant shows a trend towards higher frequency at higher voltages previously reported for EOF effects in alpha-hemolysin [40], while the wt displays an opposite tendency likely to be governed by direct rotation of the drug in the eyelet.

### Liposome-swelling shows reduced ampicillin permeation through G103K PorB

For qualitative investigation of antibiotics transport, liposome-swelling assays are often performed, in which the influx of antibiotics is measured by the decrease in optical density of bursting liposomes [6]. Liposomes containing wt and G103K PorB as well as liposomes without protein as control were monitored in the presence of ampicillin to study the drug influx through PorB (Fig. 7). The optical density (OD400) decreases over time only for the samples containing porins, showing that ampicillin is translocated into the liposomes. Ampicillin transport is substantially reduced for the mutant compared to wt PorB in these experiments, demonstrating the effect of the single mutation on overall antibiotic uptake.

**FIGURE 7.**
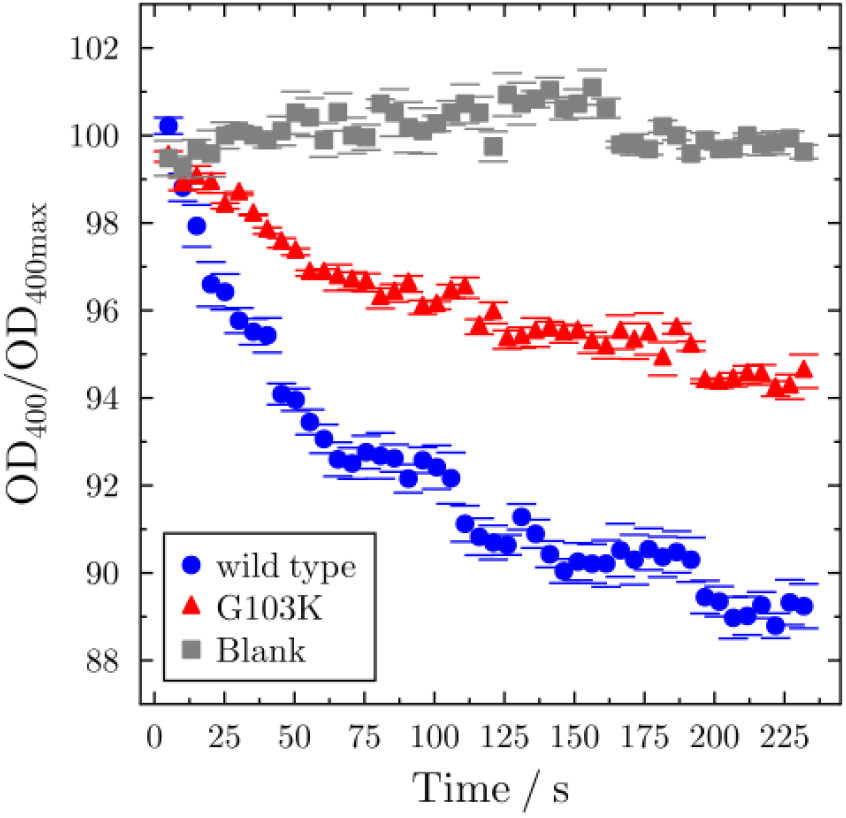
Liposome swelling assay. Liposomes containing wt PorB (blue) and G103K PorB (red) were monitored in the presence of ampicillin. Liposomes without protein (blank, gray) were used as control. The mean values of three independent measurements with the standard deviation (bars) are plotted.

## CONCLUSIONS

The eyelet of PorB, and other porins, is shaped to facilitate the selective translocation of a range of polar solutes into the bacterial cell. Its geometry and transverse electrostatic field, arising from clusters of basic and acidic residues in the eyelet, create two distinct pathways for cations and anions. The G103K mutation, related to drug-resistant phenotypes, introduces a bulkier and positively charged sidechain into the centre of the PorB eyelet. This affects the selectivity of the channel by reducing its size and altering its electrostatic properties. Despite the mobility of the lysine sidechain, a narrower pore of G103K PorB is seen in both X-ray structures and molecular dynamics simulations. The G103K mutation also changes the bipolarity of the eyelet by introducing a positive charge into the channel constriction. The structural and electrostatic effects translate into a lower overall ion conductance of this porin, as observed by high-resolution experimental electrophysiology. The decreased conductance is also manifested in our CompEL simulations, which show a lower conductance, and the potentials-of-mean-force for cations and anions along the pore axis which reveal a disruption of the cation pathway due to the additional charge in the eyelet.

Both our electrophysiology experiments and CompEl simulations show that, despite similar binding sites for ampicillin near the eyelet, ampicillin-bound wt and G103K PorB behave differently with respect to ion flux and blocking frequency under voltage (Fig. 5). Our simulations reveal a difference between the effects of voltage on the binding of ampicillin in the mutant in a different way compared to the wt. In the mutant, high voltages lead to rotation of the ligand out of its binding site, whereas in the wt, translation due to the EOF predominates. Ultimately, the difference arises due to the changed polarity and geometry of the eyelet in the mutant. To ascertain if the mutation also affects the permeability of PorB for ampicillin, we carried out liposome swelling assays. The results show that G103K PorB indeed exhibits a reduced experimental permeability for ampicillin, with an approximately two-fold reduced rate.

## METHODS

### Crystal structure and electrostatic surface potential of PorB G103K

PorB G103K from *Neisseria meningitidis* W135 strains was expressed, purified and crystallized as described previously [17, 23, 42, 43]. Crystals of PorB in the P6_3_ crystal form were obtained in 100 mM MES, pH 6.5, 31% Jeffamine M-600. Diffraction data were collected at the Swiss Light Source (SLS) PX-I (X06SA) beamline at a wavelength of 1.0 Å. Datasets were processed with HKL-2000 suite [44]. Data collection and refinement statistics are listed in Table S1 (Supporting Information). A simple refinement using previously determined crystals of PorB wild type (PDB ID: 3VY8) [24] was performed with REFMAC5 [45] in CCP4 suite [46] to obtain initial model phases. Iterative rounds of manual model building in *Coot* [47] and refinement in REFMAC5 [45] were used to improve the quality of both models. Structure validation was carried out with PROCHECK [48]. Figures were prepared using PyMOL (version 1.7.6.6, https://www.pymol.org or CueMol (version 2.2.1.354, http://www.cuemol.org/en/), and electrostatic surface potential comparisons were calculated using the Adaptive Poisson-Boltzmann Solver (APBS) [26].

The coordinates and the structure factors for the PorB mutant G103K have been deposited in the RCSB Protein Data Bank under the accession code 7DE8.

### Liposome swelling assay

To test for ampicillin transport activity of PorB wt and PorB G103K, a liposome swelling assay was performed at isoosmotic conditions as described previously [42]. Multilamellar liposomes were prepared using *E. coli* polar lipid extract (Avanti Polar Lipid, Alabaster) [21, 23].

Lipids (2 mg) were dissolved in chloroform and dried under vacuum for 20 min. Subsequently, the lipid film was rinsed with 800 µl n-pentane and resuspended in water under constant agitation for 60 min. The resulting suspension was equally split into two samples, one containing PorB, the other one serving as the control sample (1 mg lipid plus 0.5 μg PorB or an equal volume of buffer solution, respectively). Both liposome preparations were sonified until they appeared translucent (2-3 min). After drying in a speedvac for 2 h, the samples were stored in a dark desiccator overnight. Films were resuspended in 800 μl buffer solution (12 mM stachyose, 1 mM NAD-imidazole, 4 mM Na-NAD, pH 6.0) followed by incubation at room temperature, first for 1 h under gentle movement, then for 1 h without. The swelling assay was performed by addition of 10 µl of liposomes to 500 µl ampicillin solution (15 mM in 1 mM NAD-imidazole, pH 6.0) and the optical density was recorded at a wavelength of 400 nm with a resolution of 5 s. Three independent experiments were performed for each sample.

### Electrophysiological recordings on PorB

Single channel recordings of PorB were performed on solvent-free planar bilayers using the Port-a-Patch (Nanion Technologies, Munich, Germany) as described previously. Briefly, a giant unilamellar vesicle (GUV) composed of 1,2-diphytanoyl-*sn*-glycero-3-phosphocholine (DPhPC)/cholesterol (9:1) was spread in 1 M KCl, 10 mM HEPES, pH 7.5 on the aperture (*d* = 1-5 µm) in a borosilicate chip by applying 10-40 mbar negative pressure resulting in a solvent-free membrane with a resistance in the GΩ range. Afterwards, PorB stock solution (2.2 µM in 200 mM NaCl, 20 mM Tris, 0.1 % (*w*/*w*) LDAO (*N,N*-dimethyldodecylamine *N*-oxide), pH 7.5) was added to the buffer solution (50 µL) at an applied DC potential of +40 mV. Current traces were recorded at a sampling rate of 10 kHz and filtered with a low-pass four-pole Bessel filter of 1 kHz using an Axopatch 200B amplifier (Axon Instruments, Union City, CA, USA). For digitalization, an A/D converter (Digidata 1322; Axon Instruments) was used and data analysis was performed with the Clampfit 10.4.0.36 from the pClamp 10 software package (Molecular Devices, Sunnyvale, CA, USA).

For electrophysiological measurements in the presence of ampicillin, planar black lipid membranes (BLMs) were prepared by adding 1-2 µL of lipids (DPhPC/cholesterol, 9:1) dissolved in *n*-decane (30 mg/mL) to an aperture (*d* = 50 µm) in a PTFE foil (DF100 cast film, Saint-Gobain Performance Plastics, Rochdale, Great Britain) fixed between two cylindrical PTFE-chambers filled with 3.0 mL buffer (1 M KCL, 10 mM HEPES, pH 7.5 and pH 6.0, respectively). Protein was added to the *cis* chamber and inserted by stirring at an applied DC potential of +40 mV. After protein insertion, ampicillin was added from a stock solution (25 mM in 1 M KCl, 10 mM HEPES, pH 7.5 and pH 6.0, respectively) to both sides of the BLM. Current traces were recorded at a sampling rate of 50 kHz and filtered at 5 kHz. Ampicillin blocking events were analysed using JULES for model-free idealization of the data [29]. Its combination of multiresolution techniques and local deconvolution allows a precise idealization of events below the filter length, in particular amplitudes and residence times that are smoothed by the filter are reconstructed with high precision as shown previously [13]. Results are confirmed by an analysis with a hidden Markov model, in which the filter is taken into account explicitly.

### Docking

The binding mode of ampicillin was explored by means of docking calculations carried out with GOLD [35] and rDock [36–38]. Docking computations were performed with a 2-fold purpose to explore suitable starting orientations of the inhibitor in the binding site of the wildtype and G103K mutant PorB. The structural models of PorB included in the docking calculations were the X-ray structures of the PorB (wt, PDB ID 3VY8 [24] and the G103K mutant described in the present paper) and extra structures were extracted for the apo simulations and minimized. Each compound was subjected to 100 docking runs. Whereas the protein was kept rigid, GOLD and rDock account for the conformational flexibility of the ligand around rotatable bonds during docking calculations. The output docking modes were analysed by visual inspection in conjunction with the docking scores.

### Set up of the PorB Systems

The PorB wild type and the G103K variant were respectively modelled using the X-ray structure obtained by Kattner et al. (PDB ID 3VY8) [24] and the X-ray structure described in this work. The PorB trimers were embedded into a preequilibrated 150 x 150 Å 1-palmitoyl-2-oleoyl-*sn*-glycero-3-phosphocholine (POPC) bilayer. A 25 Å layer of SPC/E water molecules was set up at both sides of the bilayer, and K^+^ cations and Cl^-^ anions were added to achieve an ionic strength of 1 M to enhance the sampling of ion passage along the protein pores (approximately 57000 water molecules and 1866 K^+^ and 1896 Cl^-^ ions). PorB trimers were embedded into a lipid bilayer composed of 542 POPC lipid molecules using GROMACS utility membed [49, 50]. The Parm99SB force field [51, 52] and virtual sites for hydrogen atoms [53] were used for the protein. The POPC molecules were parameterized according to the lipid parameters derived by Berger et al. [52, 54], the SPC/E water model was used to model water molecules [55] and Joung and Cheatham parameters [56] were used to model the counterions. The zwitterionic ampicillin molecule was parameterized using the gaff force field [57] in conjunction with RESP (HF/6-31G(d)) charges [58] as implemented in the Antechamber module of AMBER12 software package [59].

### Molecular Dynamics Simulations

Molecular simulations were carried out with the GROMACS molecular dynamics package, version 5.1.5. [60]. For each system, the geometry was minimized in four cycles that combined 3500 steps of steepest descent algorithm followed by 4500 of conjugate gradient. Thermalization of the system was performed in 6 steps of 5 ns, where the temperature was gradually increased from 50 K to 320 K, while the protein was restrained with a force constant of 10 kJ mol^-1^ Å^-2^. The systems were equilibrated with restrained protein for 100 ns to ensure the equilibration of the aqueous phase. Production runs accounted of 200 ns long simulations.

The temperature was kept constant by weakly coupling (*t* = 0.1 ps) the membrane, protein, and solvent separately to a temperature bath of 320 K with the velocity-rescale thermostat of Bussi *et al*. [61]. The slightly higher temperature was chosen to ensure that the membrane remains in a fluid phase. The pressure was kept constant at 1 bar using semi-isotropic Berendsen coupling [62]. Long-range electrostatic interactions were calculated using the smooth particle mesh Ewald method [63] beyond a short-range Coulomb cut-off of 10 Å. A 10 Å cut-off was also set for Lennard-Jones interactions. LINCS algorithm [64] was used to restrain the system and SETTLE algorithm [65] was used to constrain bond lengths and angles of water molecules. Periodic boundary conditions were applied. Taking advantage of the Berger lipid model and the virtual sites, the integration time-step was set to 4 fs in the *apo* PorB simulations and 2 fs in the *holo* PorB simulations.

### PMF landscape calculations

Productions runs for each system were used to obtain two-dimensional potential-of-mean-force (PMF) free energy landscape of the passage of K^+^ and Cl^-^ ions along PorB pore taking into account data sets from all channels. The PMF was generated by binning the system along the axis of the translocation pathway *z* and along an orthogonal axis, referred to as *x*, corresponding to a vector from the Center Of Masses (COM) of the acidic cluster to the COM of the basic cluster in the eyelet region. A bin length of 0.5 Å was used in both directions.

The PMF was calculated from the ionic density *n*_*i*_(*x,z*) in each bin according to:

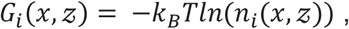

where *k*_B_ is Boltzmann’s constant and *T* is the temperature. The density of both ion types *i* within each bin was determined from the full atomic trajectory, normalized to the channel area at the pore entry. PMF values are shown as the free energy difference relative to minimum free energy for each ion data set.

## AUTHOR CONTRIBUTIONS

A.B., C.K., M.S. and M.T. performed the experiments, F.P., M.D. and A.M. developed and applied the mathematical tools to evaluate the data, C.M.I., S.L. and U.Z. performed the MD simulations. C.S. and U.Z. designed the experiments and wrote the manuscript.

## ACKNOWLEDGEMENTS

We are grateful to N. Denkert and M. Meinecke for support in constructing and performing the measurements on black lipid membranes and I. Mey for helpful discussions. Funded by the Deutsche Forschungsgemeinschaft (DFG, Research Foundation Germany)” under Germany’s Excellence Strategy – EXC 2067/1-390729940 (C.S. and A.M.), the Bundesministerium für Bildung und Forschung (BMBF) program ZIK HALOmem (FKZ 03Z2HN21 to M.T.), by Grants-in-Aids for young scientist from the MEXT (No. 16K18506 to M.T.), by the Wellcome Trust Interdisciplinary Research Funds (grant WT097818MF), the Scottish Universities’ Physics Alliance (SUPA) and the Tayside Charitable Trust (U.Z. and S.L.), and by the MRC Doctoral Training Programme (C.M.I). The coordinates and the structure factors for the PorB mutant G103K have been deposited in the RCSB Protein Data Bank under the accession code 7DE8.

