## Supplementary Information for "An antibiotic-resistance conferring mutation in a neisserial porin: Structure, ion flux, and ampicillin binding"

**LIST OF FIGURES**

**Table S1. Crystallographic data.** Data collection and refinement statistics. 3

**Figure S1.** Electrostatic surface potential of PorB viewed from the periplasmic side, calculated with APBS. 4

**Figure S2.** Electrostatic surface potential of PorB viewed from the extracellular side, calculated with APBS. 5

**Figure S3.** Event histogram of conductances recorded on a solvent-free membrane. 6

**Figure S4.** RMSD fluctuations of the unbiased Molecular dynamics simulations of apo PorB systems. 7

**Figure S5.** Schematic representation of the *x* and *z* axes used to obtain the PMF plots of the $K^+$ and $Cl^-$ ions along the PorB pore. 8

**Figure S6.** Unbiased molecular dynamics simulations of ampicillin-bound PorB systems**.** **9**

**Figure S7.** Comparison of field-dependent translocation of ampicillin in wt and G103K PorB systems. 10

**Figure S8.** Effect of the ampicillin binding on the ion current in simulations under voltage. **11**

**Figure S9.** Field-dependent rotation of ampicillin in wt PorB simulations under voltage. 12

**Figure S10.** Conductance of ampicillin-bound wt and G103K PorB trimers as a function of time. 13

**Table S1. Crystallographic data.** Data collection and refinement statistics.

| | |
|---|---|
| **Data collection** | |
| Beamline | SLS PXI (X06SA) |
| Detector | Pilatus 16M |
| Wavelength (Å) | 1.00 |
| Space group | $P6_3$ |
| **Cell dimensions** | |
| *a*, *b*, *c* (Å) | 84.39, 84.39, 107.11 |
| $\alpha$, $\beta$, $\gamma$ (°) | 90, 90, 120 |
| Resolution (Å) | 50-2.75 (2.81-2.75)[a] |
| Total no. of reflections | 75393 |
| Unique no. of reflections | 11223 |
| Redundancy | 6.7 (7.1) |
| $R_{merge}$ (%)[b] | 0.077 (0.840) |
| $I/\sigma I$ | 22.67 (2.22) |
| Completeness (%) | 99.8 (100) |
| **Refinement** | |
| Resolution (Å) | 2.76 |
| $R_{work}$[d]/$R_{free}$[e] | 21.3/26.0 |
| **No. of atoms** | |
| Protein | 2612 |
| **RMSD**[f] | |
| Bond lengths (Å) | 0.008 |
| Bond angles (⁰) | 1.466 |
| **Ramachandran plot** | |
| Favored (%) | 97.6 |
| Allowed (%) | 2.4 |
| Outliers (%) | 0 |

[a]Values in parentheses are for the highest resolution shell.

[b]$R_{merge} = \Sigma|I_{obs} - I_{avg}|/\Sigma I_{avg}$, where $I_{obs}$ is the measured intensity of each reflection, and $I_{avg}$ is the intensity averaged from symmetry equivalents.

[d]$R_{work}= (\Sigma||F_o| - |F_c|| / \Sigma|F_o|)$, where $F_o$ denotes the observed structure factor amplitude and $F_c$ denotes the structure factor amplitude calculated from the model.

[e]$R_{free}$ = $R_{free}$ is as for $R_{work}$ but calculated with 5% of randomly chosen reflections omitted from the refinement.

[f]RMSD = Root mean square deviation.

### Electrostatic surface potential of PorB

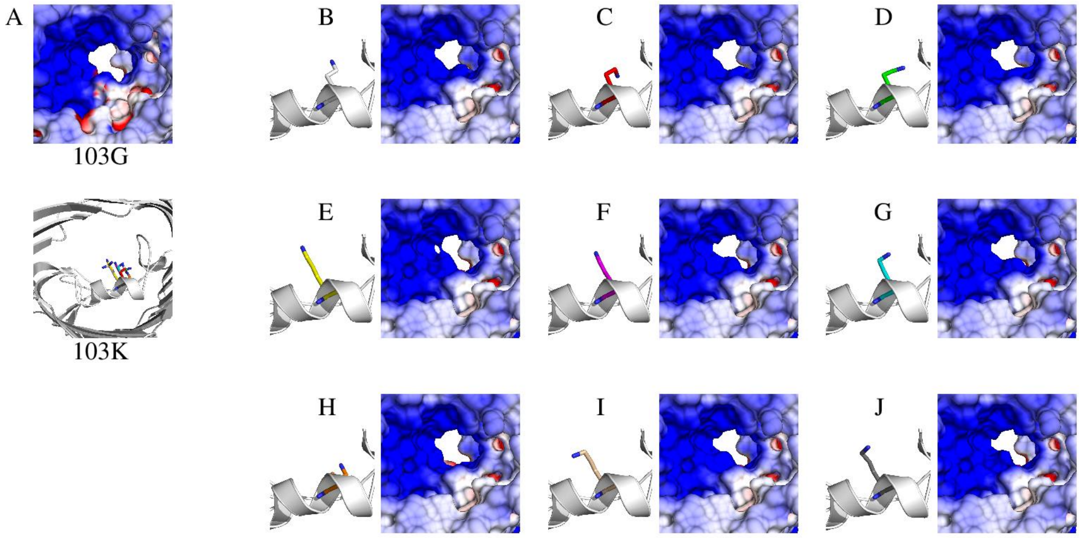


**Figure S1.** Electrostatic surface potential of PorB viewed from the periplasmic side, calculated with APBS (61). The potential of **(A)** PorB wt and **(B-J)** 9 different possible Lys103 positions (PorB G103K) are shown. Positive potential is colored in blue, and negative potential is colored in red. Potentials are contoured from +8 kT/e to −8 kT/e. The results are shown at the solvent accessible surface area of PorB.

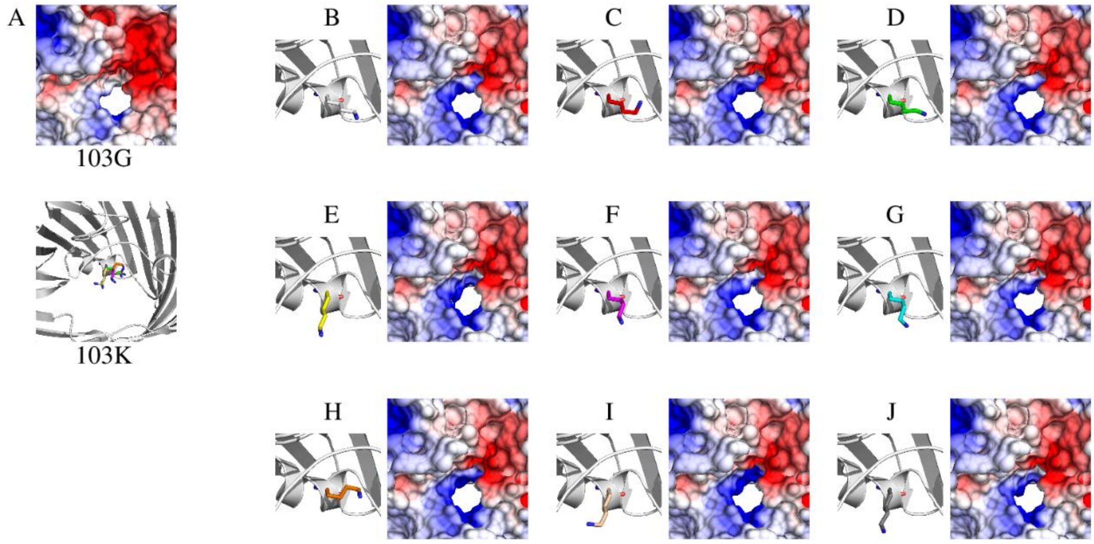


**Figure S2.** Electrostatic surface potential of PorB viewed from the extracellular side, calculated with APBS (61). The potential of **(A)** PorB wt and **(B-J)** 9 different possible Lys103 positions (PorB G103K) are shown. Positive potential is colored in blue, and negative potential is colored in red. Potentials are contoured from +8 kT⁄e to −8 kT⁄e. The results are shown at the solvent accessible surface area of PorB.

**Control recordings on solvent-free bilayers lacking PorB**

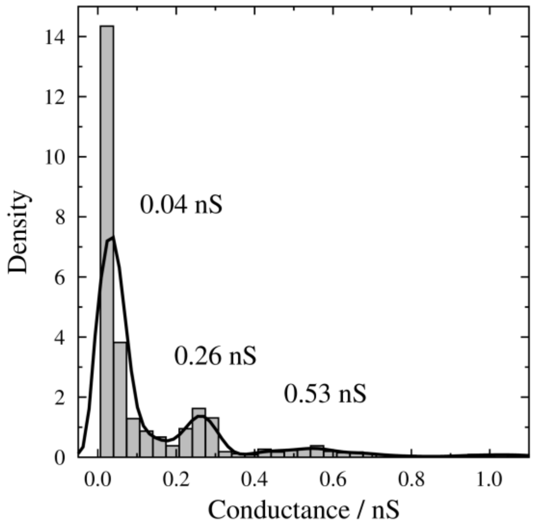


**Figure S3.** Conductivity event histogram recorded on a solvent-free membrane in 1 M KCl at pH 7.5 to which only buffer containing LDAO (200 mM NaCl, 20 mM TRIS, 0.1 % (w/w) LDAO) was added. Conductivity maxima of $G_1 = (0.04 \pm 0.02)$ nS, $G_2 = (0.26 \pm 0.07)$ nS and $G_3 = (0.53 \pm 0.16)$ nS were determined ($n$ = 2577) with a kernel density estimation using a Gaussian kernel (black solid line)

### Atomistic simulations of the ionic pathway

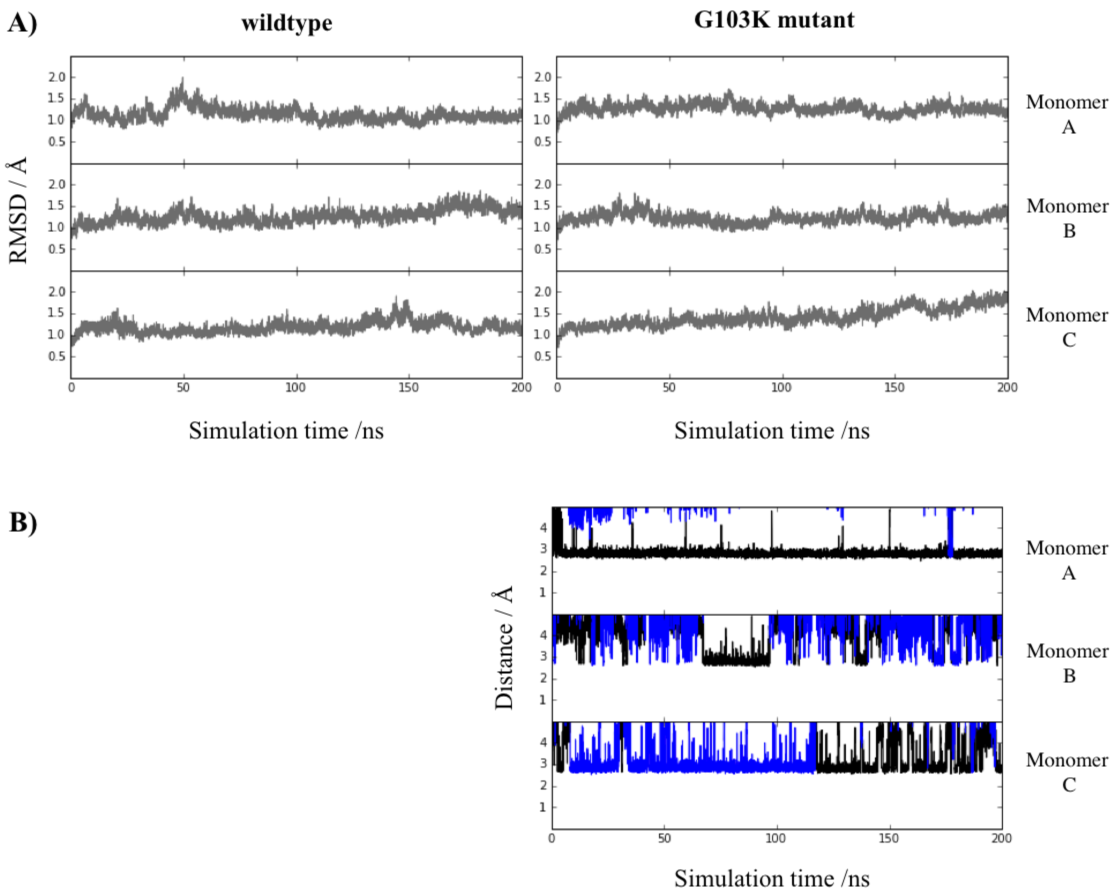


**Figure S4.** Unbiased molecular dynamics simulations of apo PorB systems. **(A)** Root mean squared deviation (RMSD) calculated for the Cα (grey) of each monomer of the wild type (left) and G103K (right) PorB simulations. **(B)** Interaction between the nitrogen atom of the Lys103 and the oxygen atoms of the residues Asp104 (blue lines) and Glu128 (black lines).

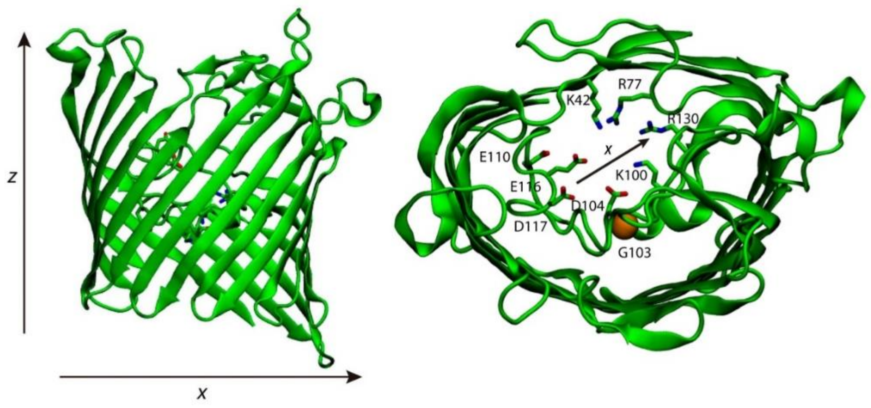


**Figure S5.** Schematic representation of the x and z axes used to obtain the PMF plots of the $K^+$ and $Cl^-$ ions along the PorB pore.

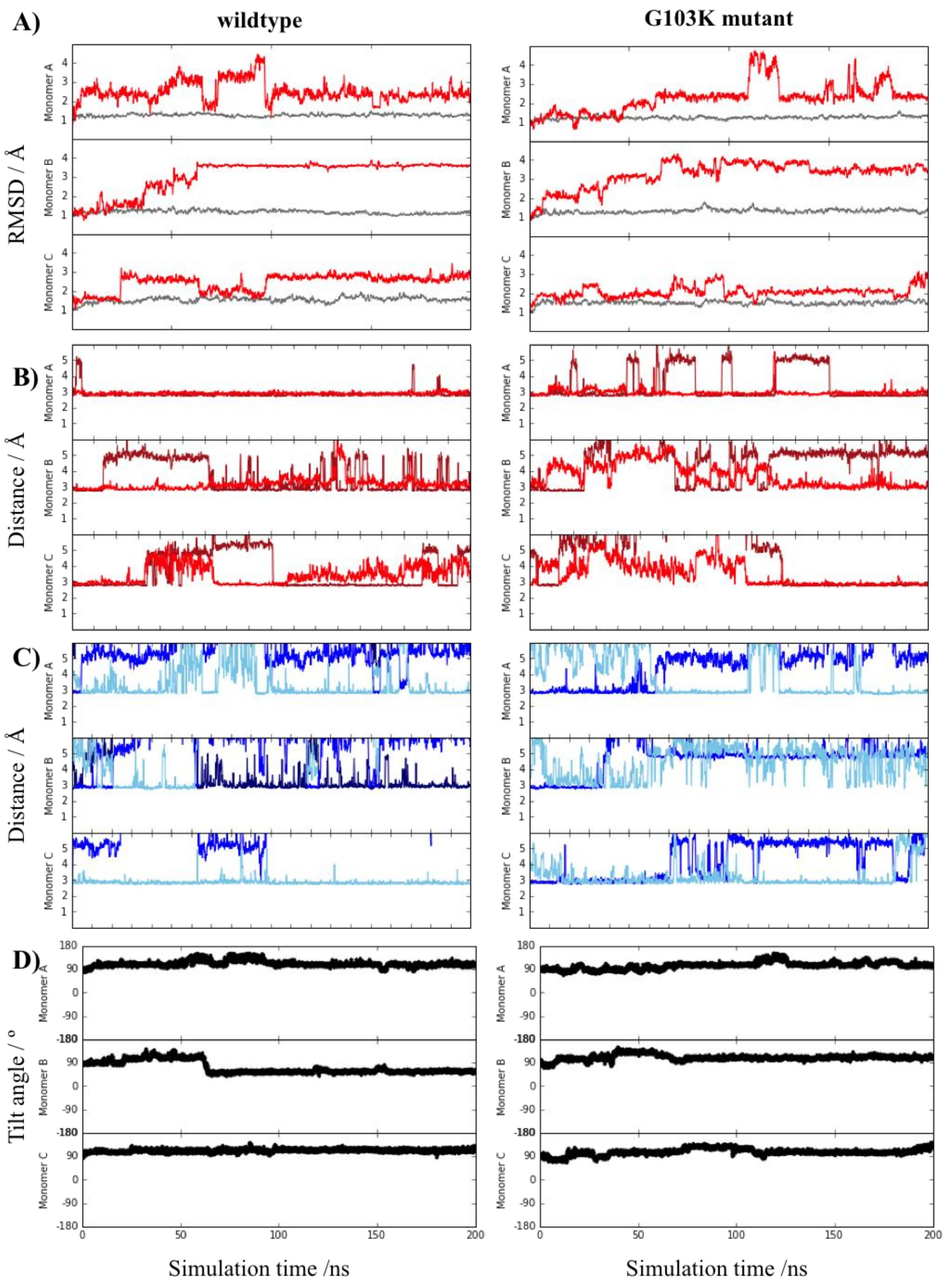


**Figure S6.** Unbiased molecular dynamics simulations of ampicillin-bound PorB systems**.** (**A**) Root mean squared deviation (RMSD) calculated for the ampicillin molecule (red) and the C$\alpha$ (grey) of each monomer (red) of the wild type (left) and G103K (right) PorB simulations. (**B**) Interactions of the ampicillin molecule with the acid side (carbonyl atom of G112 in red lines and Glu116 in maroon lines) and (**C**) basic side (Lys64 in blue lines, Lys100 in navy lines and Arg130 in blue lines) of the eyelet. (**D**) Tilt angle of ampicillin along simulation time.

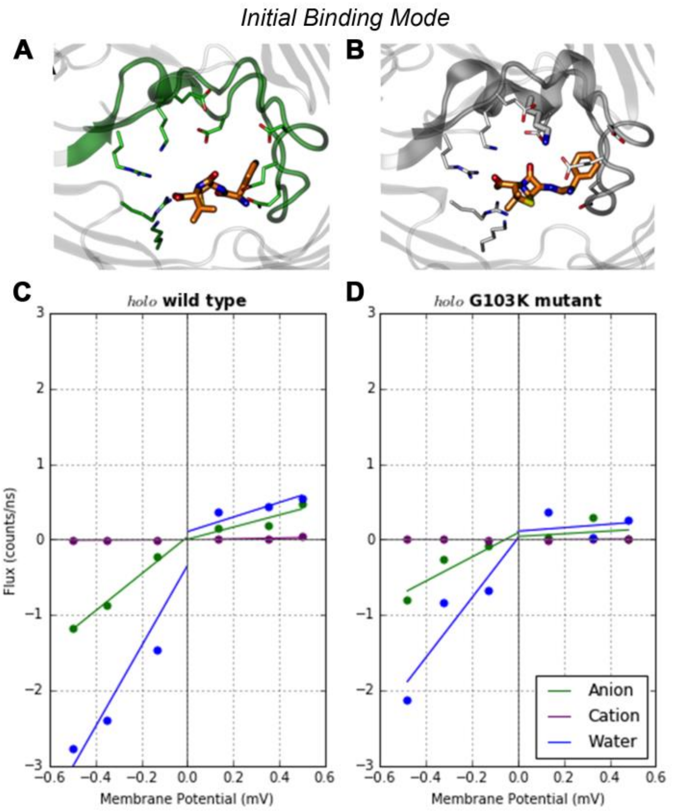


**Figure S7.** Effect of the ampicillin binding on the ion current in simulations under voltage. **(A, B)** Initial binding modes of ampicillin of the wt and G103K PorB respectively. wt and G103K PorB proteins are shown in green and grey cartoon respectively. Ampicillin molecules are shown in orange sticks. **(C,D)** The asymmetric flux of $Cl^-$ (green), $K^+$ ions (purple) and electro-osmotic flow of water molecules (blue) across the eyelet at negative and positive voltages is shown. Positive values of flux account for permeations from the extracellular compartment to the periplasmic compartment.

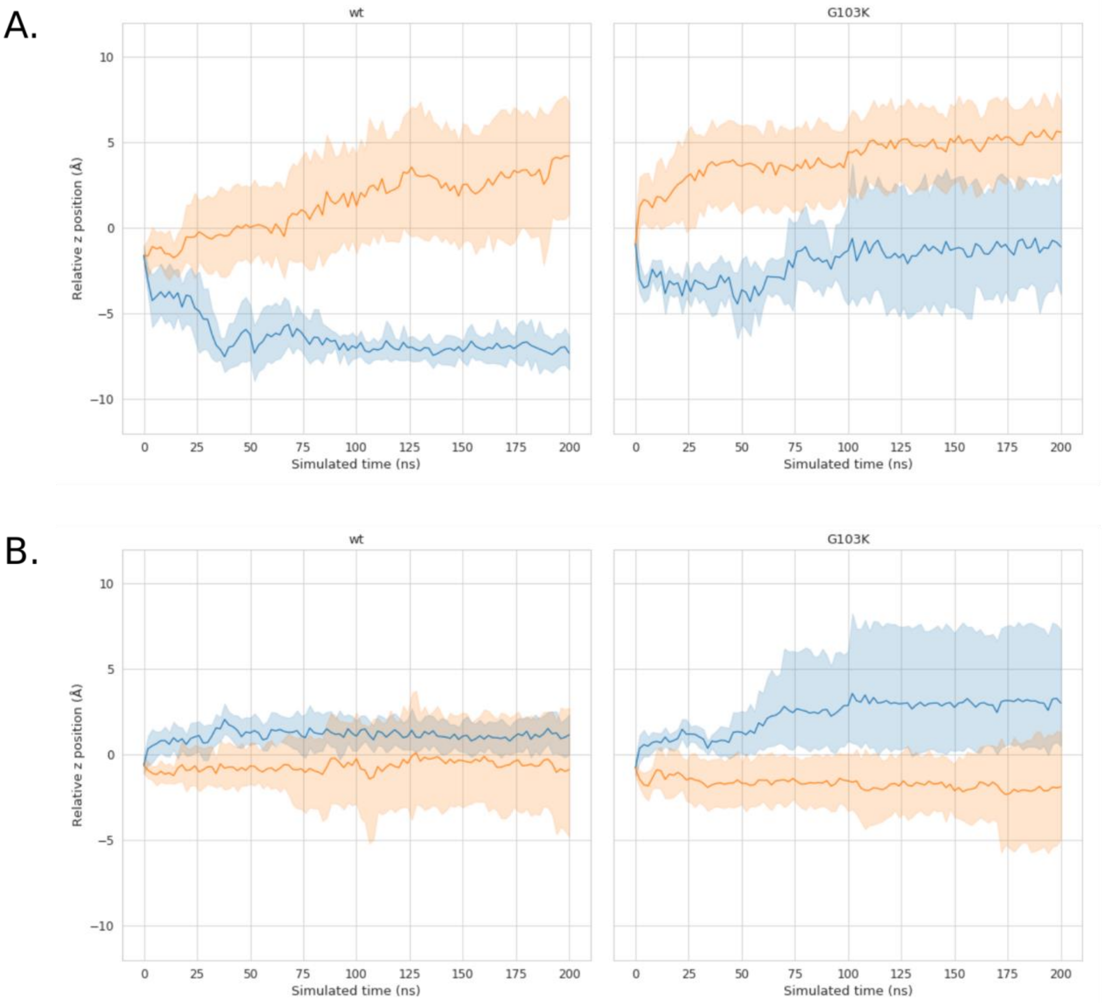


**Figure S8.** Comparison of field-dependent translocation of ampicillin in wt and G103K PorB systems. The membranes under positive and negative voltages are depicted in orange and blue, respectively, for an ion imbalance Δq of 18 (Vm ≈ ±600 mV). **(A)** Rotation of ampicillin as shown by the difference in the centre of geometry of the benzene and acidic moieties. **(B)** Translation of ampicillin under voltage in wt and G103K PorB relative to the original binding site. In the wt system, ampicillin tends to remain in the binding site but shows some rotation of the molecule. Ampicillin in G103K PorB displays a smaller degree of rotation but is translated farther from the original binding site under negative voltages. In both plots, each channel is treated individually with mean value and a confidence interval of 95% plotted every 2 ns.

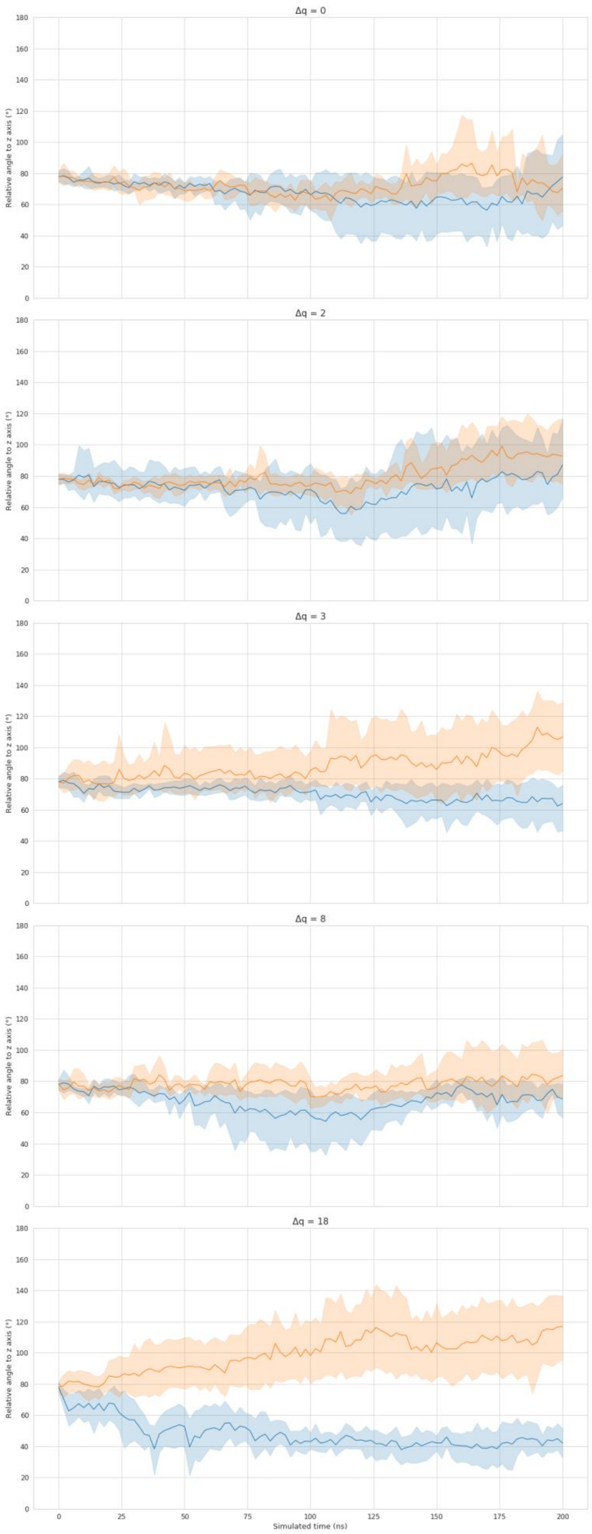


**Figure S9.** Field-dependent rotation of ampicillin in wt PorB simulations. Duplicate simulations were performed for each ion imbalance, with each pore being treated separately, resulting in data from six pores for both positive (orange) and negative (blue) voltages. The angle of the drug to the z-axis is defined as the angle formed between the z-angle and a plane between the benzene and carboxylic moieties. The mean angle is plotted every 2 ns with a confidence interval of 95%. The data shown are for ion imbalances in the CompEL double membrane system of 0, 2, 3, 8 and 18e from top to bottom.

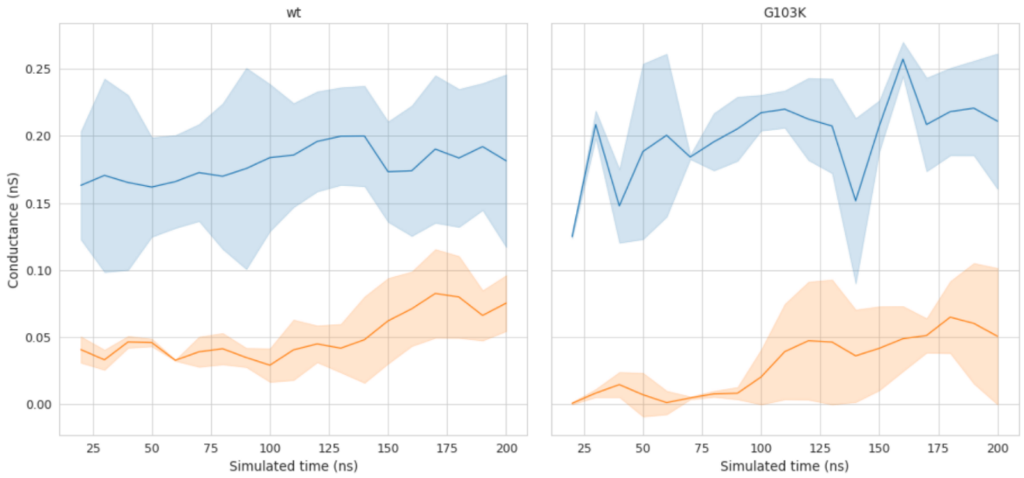


**Figure S10.** Conductance of ampicillin-bound wt and G103K PorB trimers as a function of time. The membranes under positive and negative voltages are depicted in orange and blue, respectively, for a CompEL ion imbalance Δq of 18e (Vm ≈ ±600 mV). The voltage and number of ion permeations were calculated within overlapping 20 ns windows. The conductance refers to the homotrimer system, with the mean value of each duplicate system plotted with a confidence interval of 95%.
